# AMAW: automated gene annotation for non-model eukaryotic genomes

**DOI:** 10.1101/2021.12.07.471566

**Authors:** Loïc Meunier, Denis Baurain, Luc Cornet

**Affiliations:** InBioS-PhytoSYSTEMS, Eukaryotic Phylogenomics, University of Liège, B-4000 Liège, Belgium; TERRA Teaching and Research Centre, Microbial Processes and Interactions, Gembloux Agro-Bio Tech, University of Liège, B-5030 Gembloux, Belgium; BCCM/IHEM, Mycology and Aerobiology, Sciensano, B-1000 Bruxelles, Belgium

## Abstract

**Summary:** To support small and large-scale genome annotation projects, we present AMAW (Automated MAKER2 Annotation Wrapper), a program devised to annotate non-model unicellular eukaryotic genomes by automating the acquisition of evidence data (transcripts and proteins) and facilitating the use of MAKER2, a widely adopted software suite for the annotation of eukaryotic genomes. Moreover, AMAW exists as a Singularity container recipe easy to deploy on a grid computer, thereby overcoming the tricky installation of MAKER2.

**Availability:** AMAW is released both as a Singularity container recipe and a standalone Perl script (https://bitbucket.org/phylogeno/amaw/).

**Supplementary information:** Supplementary data are available at *Bioinformatics* online.

## 1 Introduction

Coding sequences (CDS) from an organism are essential genomic data, especially for phylogenomics and gene mining, in which obtaining accurate protein sequences from publicly available emerging draft genomes is invaluable. These CDS can be obtained by gene prediction or by the structural annotation of a genome, a more complete process aiming to define the whole gene structure, which includes UTRs (UnTranslated Regions) (Yandell and Ence, 2012).

Following the decrease in sequencing costs due to the advent of NGS and the concomitant explosion of sequenced organisms, new genomic data from emerging model organisms allow researchers to access unexplored lineages, so as to expand our knowledge of poorly represented taxonomic groups. However, eukaryotic genomes, whose biodiversity is predominantly represented by protist lineages (Adl *et al*., 2019, Burki *et al*., 2020), present special features (i.e., large genomes with low gene density - long intergenic regions - as well as introns), which complexify the structural annotation process (Yandell and Ence, 2012). Although pipelines for eukaryotic genome annotation have been developed for more than a decade, it is still challenging to obtain an accurate annotation of the gene structure, a shortcoming that is often revealed in phylogenomic studies (Di Franco *et al*., 2019). MAKER2 (Holt and Yandell, 2011) is currently one of the most efficient annotation pipelines for eukaryotes. To address this issue, it combines different gene prediction tools (i.e., AUGUSTUS, SNAP) and takes advantage of experimental evidence data (transcripts and proteins) to limit the number of false positive *ab initio* gene predictions. Although MAKER2 enables individual laboratories to annotate non-model organisms (for which pre-existing gene models are not available), the use of this tool remains complex, as it implies the orchestration and fine-tuning of a several step process. First, an evidence dataset must be compiled by collecting phylogenetically related proteins and species-specific transcripts, which often requires the assembly of RNA-Seq data for new organisms. Next, iterative runs of MAKER2 must also be coordinated to aim for accurate predictions, which includes intermediary specific training of different gene predictor models. Here we present AMAW (Automated MAKER2 Annotation Wrapper), a wrapper pipeline facilitating the annotation of emerging unicellular eukaryotes (i.e., protist) genomes in both small and large-scale projects by efficiently orchestrating these different steps in a grid-computing environment. This tool addresses all the above-mentioned tasks according to MAKER2 authors’ recommendations (Campbell *et al*., 2015) and is, to our knowledge, the first implementation automating the use of MAKER2. We also demonstrate that the use of AMAW yields genome annotation significantly improved in comparison to the use of MAKER2 with the AUGUSTUS gene models that are available by default.

## 2 Input data

The most basic inputs required by AMAW is a FASTA-formatted nucleotide genome file and the organism name, which is used for acquiring protein and transcript data from public databases. Alternatively, evidence data, such as proteins or transcripts/ESTs provided by the user, or even gene models, can also be directly used for genome annotation.

## 3 Functionalities

The MAKER2 annotation suite was chosen to be automated for its performance and interesting features: beside supporting *ab initio* gene prediction with evidence data, MAKER2 has been demonstrated to improve the accuracy of the internal gene predictors, to maintain this accuracy even when the quality or size of evidence data decreases, as well as to limit the number of overpredictions, thus yielding a more consistent number of genes (Holt *et al*., 2011).

Taking MAKER2 as its internal engine, AMAW is able to gather and assemble mRNA-seq evidence, collect protein evidence, iteratively train the HMM models of the gene predictors, in order to yield the most accurate evidence-supported annotation possible without manual curation nor prior expertise of the organism to annotate (see Supplementary information).

Our automated annotation tool for non-model unicellular eukaryotic genomes, based on MAKER2, presents helpful applications in the phylogenomic and comparative genomic fields. Indeed, some taxonomic lineages still lack high-quality genomic data (Burki et al., 2020), and filling these gaps would extend studies to these interesting groups.

## 4 Application example: Structural genome annotation of selected organisms across distributed protist lineages

The efficiency of MAKER2 being already well known (Holt *et al*., 2011), we chose to illustrate the performance of AMAW by comparing the MAKER2-based annotation of a selection of 32 protist genomes in two very contrasted conditions. In more details, the annotations generated with AMAW, where a gene model is specifically created for the genome from the available data, are compared with those produced with gene models directly available in the AUGUSTUS library (Stanke *et al*., 2008). The latter condition corresponds to a basic usage of MAKER2, requiring no advanced bioinformatics manipulation.

For exploring the impact of the gene model choice, four AUGUSTUS models were used against the AMAW generated ones: *Homo sapiens, Arabidopsis thaliana, Aspergillus oryzae* and the “closest” available model with respect to the organism to annnotate. For this, a dataset of 32 genomes of protist organisms was designed and the quality of the different structural annotations was then assessed using the completeness metrics provided by BUSCO v4 (Seppey *et al*., 2019) and the latest orthologous databases (Kriventseva *et al*., 2018) (see Supplementary Figure 1 describing the complete results and Supplementary Table 1 for more details on the genomes, the closest gene model for each organism and the orthologous database used with each).

The analysis of median values of BUSCO metrics shows that AMAW gene models significantly improve the quality of MAKER2 annotations (Figure 1A), with a median completeness of 93% (the “closest” gene model is the second most complete with a median of 68.7%), a median rate of fragmented annotations of 3.4% (second: closest AUGUSTUS gene model with 8.2%) and a median rate of missing annotations of 4.7% (second: closest AUGUSTUS gene model with 14%). Moreover, among the five gene models assayed for each genome (see Supplemental Figure 1), AMAW performed best, giving the most complete annotation in 65.6% of cases (the second being *A. oryzae* with 18.8%), the least fragmented annotations in 40.6% of cases (the second being *A. oryzae* with 28.1%) and the lowest proportion of missing proteins in 59.4% of cases (the second being *A. thaliana* and the closest AUGUSTUS gene model, both with 12.5%) (Figure 1B). In conclusion, the use of AMAW significantly improves on average the genome annotation in comparison with a basic usage of MAKER2. Therefore, in combination with its drastically simplified installation and running of the pipeline in a grid-computing environment, AMAW stands out as an efficient tool for supporting small and large-scale genome annotation projects.

**Figure 1:**
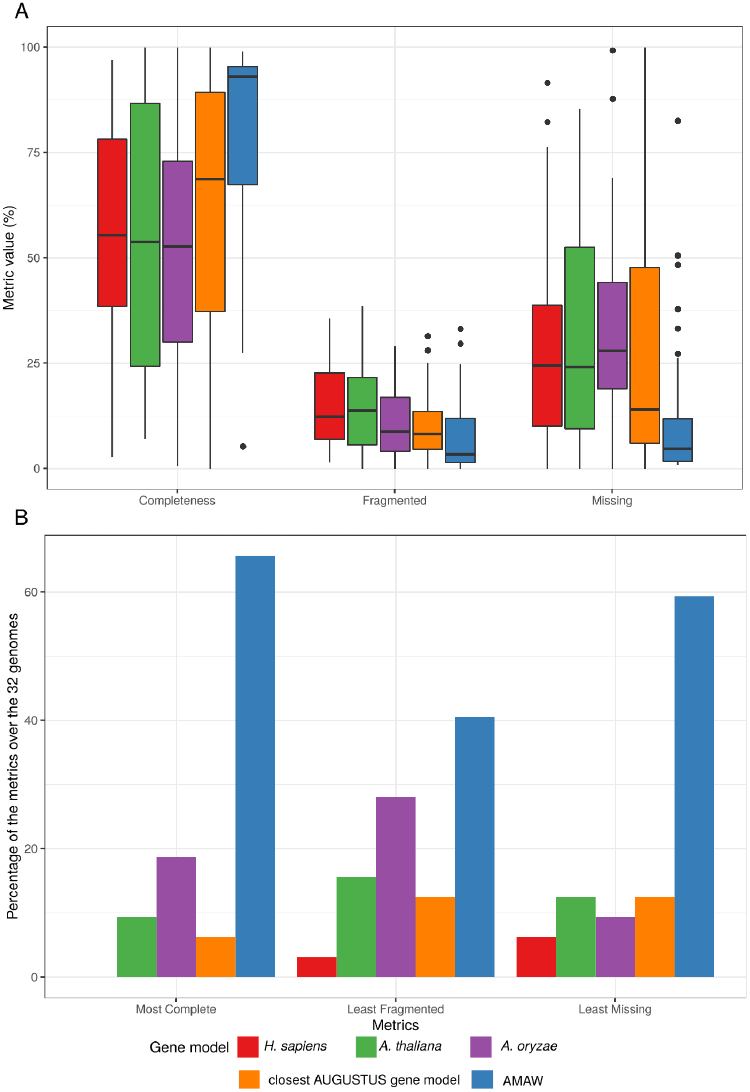
A. Median value for each gene model of the percentage of completeness, and fragmented and missing genes. B. Representation of the percentage of occurrences (out of 32 genomes) where a gene model yields the most complete annotation, the least fragmented proteins or the least missing proportion of expected proteins, in comparison with other gene models.

## Acknowledgements

We thank David Colignon (ULiège) and Olivier Mattelaer (UCLouvain) for their help with the CÉCI computing clusters.

## Supplementary information

### 1. Aim and features

The pipeline devised in AMAW aims to reach three goals: (1) to achieve the most accurate annotation of a non-model genome without manual curation, (2) to automate the use of MAKER2 for supporting large-scale annotation projects, and (3) to simplify its installation and usage for users without a strong bioinformatics background.

First, a key factor for achieving accurate genome annotation is to collect as much evidence data (ESTs, transcripts and/or proteins) as possible. This is needed both to optimize the training of specific gene models of *ab initio* gene predictors and to improve the confidence level in predictions supported by experimental data (Holt *et al*., 2011).

Second, building evidence datasets is a time-consuming task, which also implies a certain level of bioinformatics skills. Indeed, this consists, in the best cases, to find and download directly available EST, transcript or protein datasets for the genome species to annotate. However this process often further requires assembling RNA-Seq datasets into transcripts and gathering a reasonably sized protein dataset, usually including sequences of taxa phylogenetically close to the organism of interest. If building evidence datasets is feasible for a few genomes to annotate, doing so repeatedly for dozens or hundreds of genomes is hardly conceivable. This is why AMAW addresses this issue by automating the acquisition of both available RNA-Seq and protein data from reliable public databases (“NCBI SRA” for RNA-Seq data and “Ensembl genomes” for protein sequences).

Third, in addition of constructing a good input dataset for the annotation, AMAW automates the installation and the global use of the MAKER2 annotation pipeline based on good practices published by its authors (Campbell *et al*., 2015), and orchestrates the successive runs in a grid-computing environment. Even if MAKER2 is described as an easy to use pipeline, its handling and the optimal fine-tuning of its parameters demand to take notice of its large documentation and, again, require the user to have a good bioinformatics understanding.

The complete workflow of AMAW can be summarized in three steps:

1. Transcript evidence data acquisition: RNA-Seq acquisition, assembly into transcripts, quantification of the abundance of the transcripts and filtering of redundant transcripts and minor isoforms;
2. Protein evidence deployment;
3. MAKER2 Iterative runs and intermediate training of its internal gene predictors.

It is possible for the user to provide her/his in-house protein and/or transcript dataset. Moreover, one can short-circuit the pipeline by choosing an existing gene model for AUGUSTUS and/or SNAP. However, unless available models are well-suited for the organism at hand (species), it is advised to rely on AMAW full analysis.

### 2. Materials & methods

#### 2.1 Acquisition and building of transcript evidence data

The generation of a specific transcript dataset is done on the basis of the organism species name, provided by the user. This name will be used to download RNA-Seq from NCBI SRA (https://www.ncbi.nlm.nih.gov/sra). The acquisition of the RNA-Seq data prioritizes paired-end reads, when available, rather than single-end libraries for getting more accurate transcript assemblies.

FASTQ read files are assembled into transcripts with Trinity v2.12.0 (Grabherr *et al*., 2013) (standard parameters). The abundance of transcripts is first assessed with “align_and_estimate_abundance.pl”, a Trinity utilitary script that uses RSEM (Li and Dewey, 2011), then a custom script removes the redundant transcripts (which are common when several samples are pooled) and minor isoforms (by default, with abundance < 10% for a Trinity-defined gene). Finally, assembled transcripts are pooled and fetched to MAKER2.

#### 2.2 Deployment of preloaded protein evidence data

To collect a set of curated protein sequences of eukaryotic microorganisms, Ensembl genomes (Kersey *et al*., 2018) were downloaded (Protists, Fungi and Plants) and combined into a single database. However, to accelerate the computation time of MAKER2 annotations, this protein sequence database was subdivided following the major eukaryotic taxonomic clades. For this, we used the NCBI third taxonomic level (usually the phylum), which allows us to already considerably reduce the quantity of data to deploy for an annotation while ensuring enough sequence evidence for less studied lineages. Moreover, for further optimizing the computation time, these subsets were also dereplicated with CD-HIT version 4.6 (Li and Godzik, 2006): sequences sharing ≥ 99% identity were removed in favor of a single representative sequence. In practice, the taxonomy of the user-given organism species name is used to deploy the protein database corresponding to its taxon.

#### 2.3 MAKER2 runs and intermediate trainings of the *ab initio* gene predictors

Following the good practices given by Campbell *et al*. (2015), the default AMAW workflow consists in three successive MAKER2 runs:

1. The first MAKER2 round predicts the genes only based on alignment of the provided transcript and protein data on the genome assembly to annotate. The predicted gene sequences will then be used for training a gene model for the SNAP gene predictor.
2. MAKER2 second round uses SNAP with the trained gene model and the evidence data will only be used for supporting the presence or absence of the predicted genes. Then, the SNAP gene model is trained again and a gene model is trained for AUGUSTUS.
3. MAKER2 third and last round performs gene predictions with both trained SNAP and AUGUSTUS *ab initio* gene predictors.

At the end of these three annotation rounds, two sets of gene predictions containing the gene predictors consensus are returned: a first one containing those supported by evidence data and a second one with the unsupported ones. However, the latter dataset needs to be cautiously used as the false positive rate is expected to be higher.

For optimal performance of the pipeline, it is possible (and recommended when applicable) for the user to provide her/his own experimental transcript data.

Beside the complete pipeline, AMAW also offers the possibility to shorten the analyses to only one round to:

- annotate several genomes of the same species (or re-run a previous analysis) for which the evidence data has already been constructed and the SNAP and AUGUSTUS gene models already trained.
- directly use an AUGUSTUS gene model (available in its library or provided by the user) without evidence data building. It is noteworthy that this mode does not use the SNAP gene predictor.

In this case, only the third round is launched according to the chosen mode.

### 3. Assessment of AMAW performance

To assess the performance of AMAW gene models on the annotation of non-model protist organisms through the use of MAKER2, 32 genomes were selected (see Table 1 for more details on these). At the time of running, no complete proteome was available in public databases for any of these organisms (or even species of the same genera), ensuring that gene annotation was a non-trivial task for these genomes. Completeness, proportions of fragmented and missing genes for each genome annotation were computed with BUSCO v4 (Seppey et al., 2019) using the latest orthologous databases (Kriventseva et al., 2018).

The following command was used with AMAW:

~~~
singularity exec --bind <path>:/mnt annotation-amaw.sif amaw.pl \
   --genome=<genome_path> \
   --organism=<Genus_species> \
   --AUGUSTUS-db=<AUGUSTUS-config_path> \
   --AUGUSTUS-gm=<gene_model>
~~~

**Table 1:**
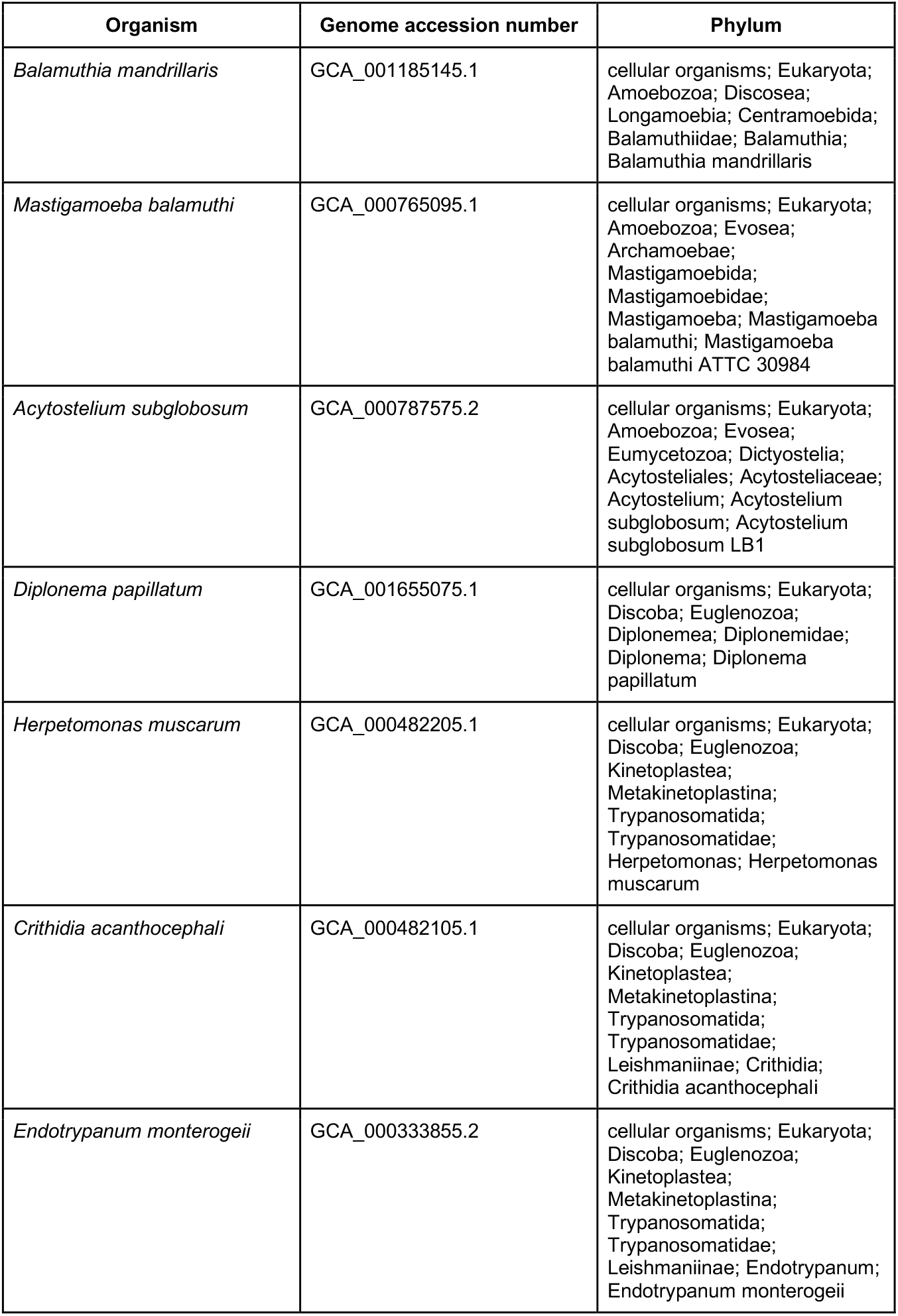
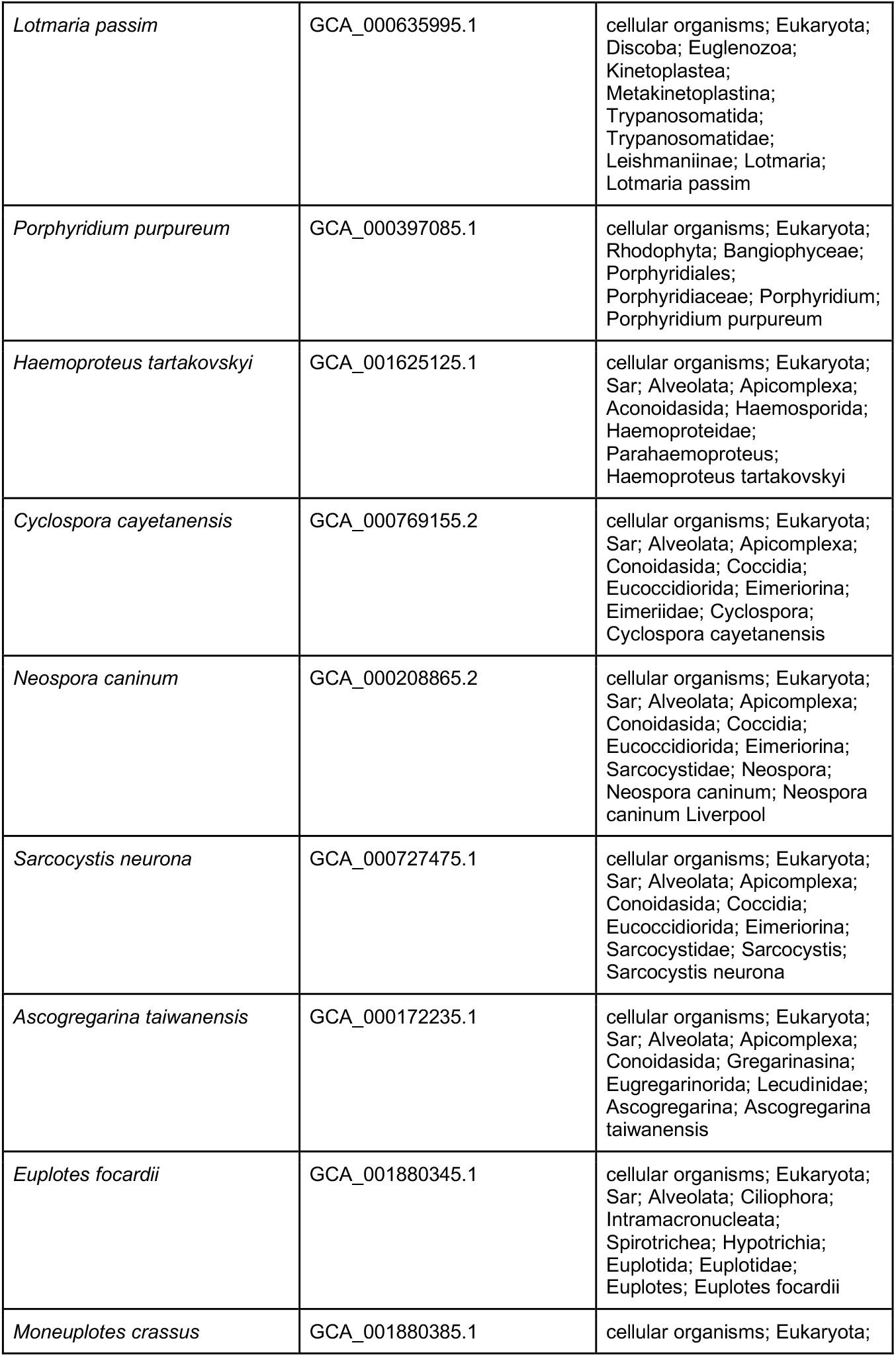
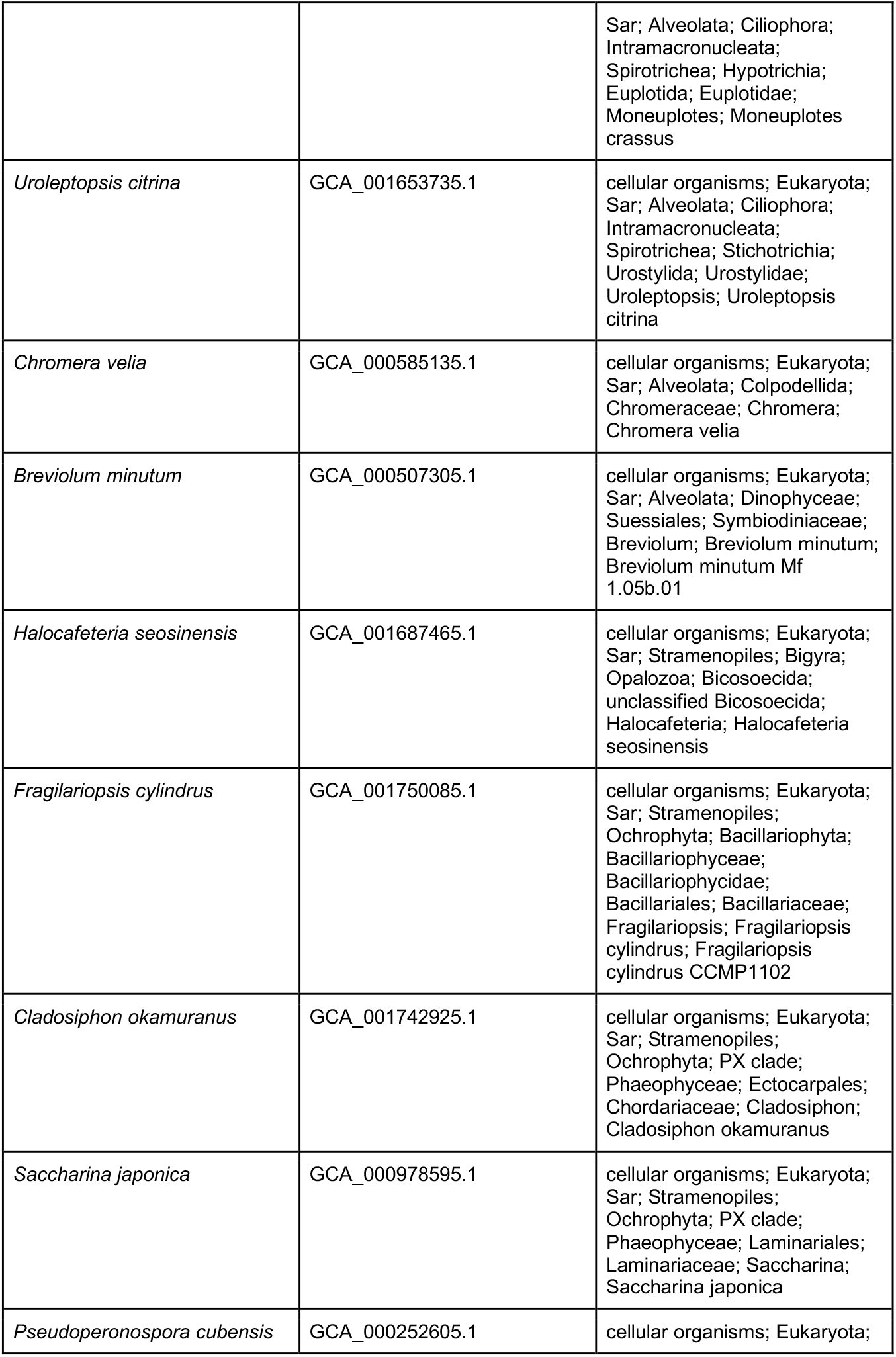
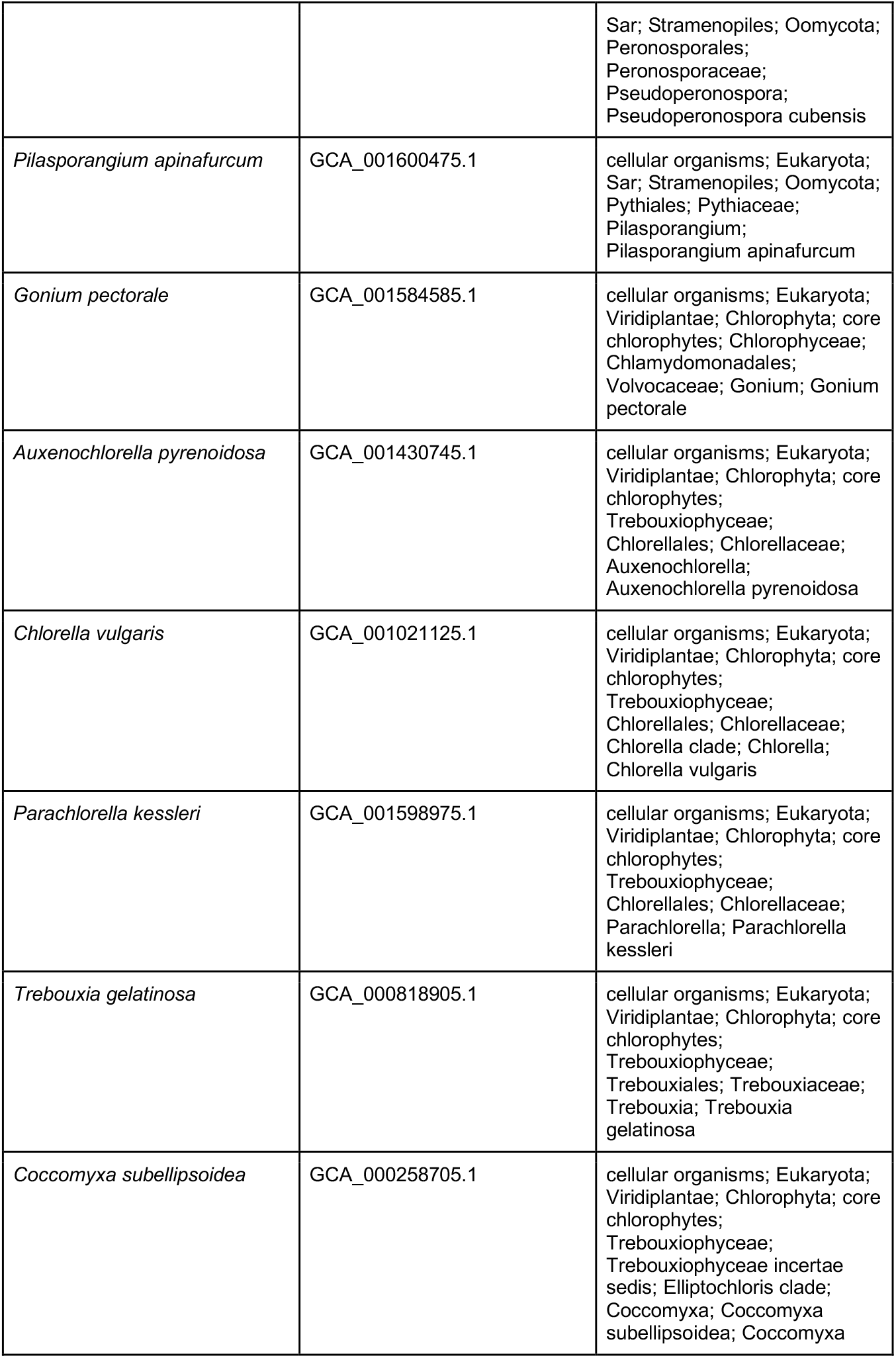
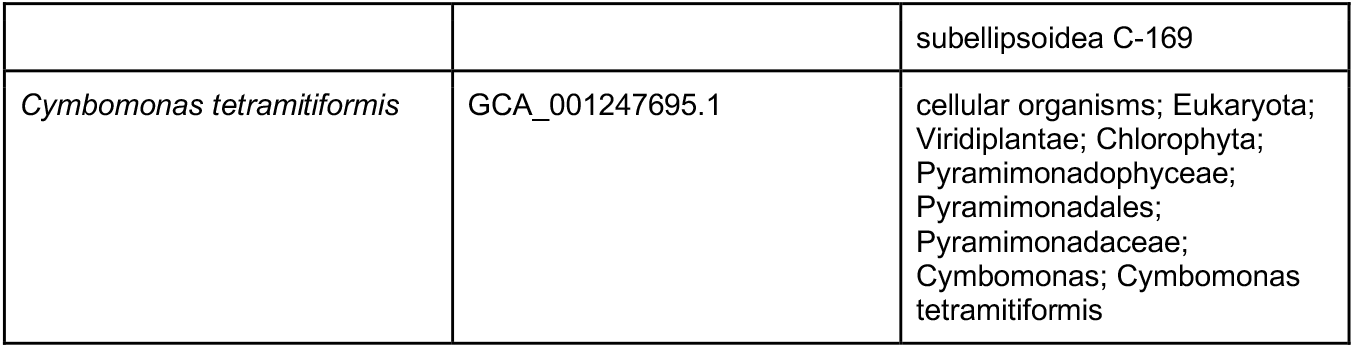
List of selected protist genomes and associated information. Organisms are sorted by taxonomic lineage, in the third column.

**Figure 1:**
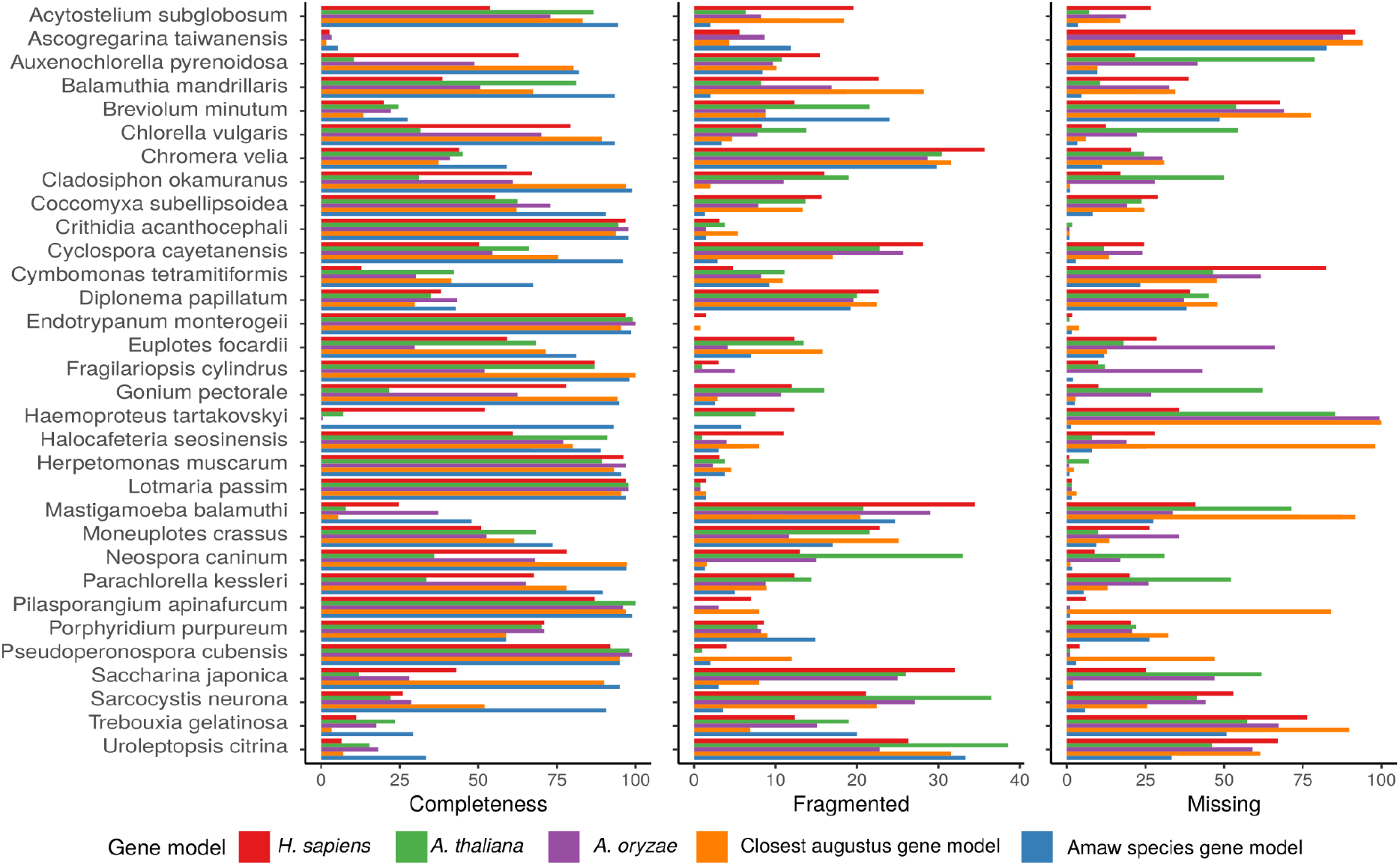
BUSCO metrics (percentage of completeness, and fragmented and missing genes) for each of the 32 analyzed genomes using five gene models. While it is desirable for a genome annotation to be the most complete, it should also be the least fragmented and having the lowest proportion of missing annotations.

